# Sphingosine 1-phosphate lyase expressed in pulmonary epithelial cells potentiates host innate defenses and alleviates influenza pathogenicity in mice

**DOI:** 10.64898/2026.07.02.736172

**Authors:** Kwang Il Jung, Savannah McKenna, Lei Jiang, Hailey Huerter, Ying He, Dong Xu, Julie D. Saba, Bumsuk Hahm

## Abstract

Influenza viruses circulate in humans, causing a substantial burden on global health. Investigation of influenza-host interactions could identify host factors that regulate influenza pathogenicity. Sphingosine 1-phosphate (S1P) is a bioactive lipid mediator and regulates crucial cellular processes. S1P lyase (SPL), an enzyme that mediates S1P degradation, was shown to display anti-influenza activity in a cell culture system. Here, we constructed a mouse model to demonstrate the antiviral function of SPL in respiratory epithelial cells during influenza in vivo. Deletion of SPL from lung epithelial cells exacerbated influenza-induced weight loss and mortality. Influenza virus began to propagate more effectively in the absence of SPL at the innate immune stage. Increased virus titers were sustained during influenza and associated with enhanced accumulation of multiple immune cell types in the lungs. Single-cell RNA sequencing was conducted to further define the function of SPL in lung epithelial cells. SPL deletion increased the proportion of alveolar type 1 (AT1) cells compared to alveolar type 2 (AT2) cells with alteration of the related signaling pathways, suggesting a role of SPL in AT1/AT2 programming. Importantly, host innate defense pathways were changed in SPL-deficient lung epithelial cells upon infection, which corroborates the antiviral function of SPL. This study elucidates the host protective function of SPL in lung epithelial cells during influenza and provides gene signature profiles critical for SPL-mediated alleviation of influenza pathogenicity. The findings may contribute to development of host-directed therapeutics to better control influenza.

## Introduction

Influenza is a continuous public health concern contributing to approximately one billion illnesses and up to an estimated 650,000 deaths globally each year^1^. Although both influenza A and B viruses circulate seasonally, only influenza A virus (IAV) has been responsible for multiple pandemics that have led to severe morbidity and mortality^2,3^. Prevention and treatment against IAV is limited to vaccinations and select antivirals, such as oseltamivir, peramivir, and baloxavir^4^. These therapeutics are limited by the tendency of the virus to undergo mutations resulting in antigenic drift, as well as its ability to undergo reassortment resulting in antigenic shift, which can lead to resistance and variable efficacy^5–8^. As such, exploration of additional and alternative therapies is warranted. Identifying novel host targets that could confer broad protection to influenza viruses would be one of the ideal approaches to combatting this constantly changing pathogen.

Sphingolipids are a family of bioactive lipid molecules that harbor a wide range of biological and immunological functions^9^. They are highly conserved and expressed in most tissues, making them an interesting target for host-directed antiviral therapeutics^10^. Previous work has suggested that the sphingolipid system plays vital roles during viral replication and undergoes metabolic shifts during viral infection^11–13^. During influenza virus infection, accumulation of specific sphingolipid species, such as ceramide and sphingosine, has been observed^14,15^. Furthermore, studies have shown a suppressive effect of ceramide on influenza virus infection, while sphingomyelin promotes influenza virus infection^16–18^. Enzymes, such as sphingosine kinases, were previously shown to be proviral during influenza virus infection, promoting production of infectious influenza viruses with enhanced viral pathogenicity^19,20^. With increasing evidence supporting influenza virus-sphingolipid interactions, more investigation into various steps of sphingolipid metabolism during viral infection is paramount to understanding the therapeutic potential of targeting this system.

Sphingosine 1-phosphate (S1P) lyase (SPL) is a sphingolipid-metabolizing enzyme that irreversibly catalyzes the cleavage of S1P into hexadecenal and phosphoethanolamine^21^. S1P is a sphingolipid that is widely recognized as a regulator of immune cell trafficking due to the chemotactic S1P gradient^22,23^. The immunosuppressive drug FTY720 (Fingolimod) is a sphingosine analog that works by downregulating an S1P receptor, sequestering lymphocytes in the lymph nodes, and is FDA approved as a treatment for multiple sclerosis^24,25^. SPL carries out the final step of S1P degradation and has been demonstrated to be critical for life^21,26^. Particularly, global deletion of SPL in mouse models leads to premature death. Defects have also been acquired in worms and fruit flies when SPL is disrupted^27^. Previous studies have shown that overexpression of SPL resulted in enhanced sensitivity to anticancer drugs in vitro^28^. In humans, activity-disrupting mutations have led to the identification of SPL insufficiency syndrome (SPLIS)^29^. Symptoms of SPLIS can range from birth defects, neurological complications, immune deficiency, endocrine disorders, and nephrotic syndrome, among others, to premature death^30–33^.

Previous work investigating SPL in the context of influenza virus infection has found that SPL harbors antiviral activity. Specifically, overexpression of SPL was found to protect infected cells, making them more resistant to virus infection^34^. Additionally, SPL was shown to promote a type I interferon (IFN) response during influenza virus infection^35^. On the other hand, it was identified that influenza virus is able to target SPL for degradation via the viral NS1 protein, dampening the antiviral effect of SPL during infection^36^. These studies provide strong evidence that SPL has antiviral therapeutic potential in the context of influenza virus infection, however, these studies were limited to in vitro cell line-based systems. This study aimed to further investigate the role of SPL in lung epithelial cells in vivo during IAV infection using an epithelial cell-specific, conditional SPL knockout (KO) mouse. We report that deletion of SPL in lung epithelial cells leads to higher virus-induced morbidity and mortality, which is associated with higher lung viral titers initiated at the innate immune stage. Single-cell RNA sequencing (scRNA-seq) analysis indicates a baseline increase in alveolar type 1 (AT1) epithelial cells in the SPL KO mouse model. Further, the gene signature pathway analysis reveals a downregulation of antiviral innate immune response and response to type I IFN, along with an upregulation in stress response signaling and apoptotic process in the absence of SPL, lending further support to an antiviral role of SPL during IAV infection in vivo.

## Results

### Epithelial-cell specific SPL knockout (SPL KO^Epi^) mice were generated to study the role of SPL in lung epithelial cells in vivo

The role of SPL has not been widely studied during infection, partly due to the lack of reliable animal models. Mice with complete deletion of SPL die prematurely due to significant complications during development^37^. To address this, the SPL floxed (SPL*^fl^*) mouse was developed and utilized to understand the role of SPL in multiple cell types^38^. This prompted us to generate a mouse model to define the role of SPL during influenza virus infection. For this purpose, we established an epithelial-cell specific SPL KO (referred to as SPL KO^Epi^) mouse as follows: SPL*^fl^* mice were crossed with Shh*^Cre+/-^*mice for four generations to produce SPL*^fl^*-Shh*^Cre+/-^* mice. Backcrossing male Shh*^Cre+/-^*with female SPL*^fl^* allowed for the generation of SPL KO^Epi^ (SPL*^fl^*-Shh*^Cre+/-^*) and littermate wild type (WT) control mice (SPL*^fl^*-Shh*^Cre-/-^*: referred to as SPL WT) to be used for experiments (Fig. 1A). Use of littermates from the same breeding colony as control mice minimizes variable factors, such as background and microbiome, and increases scientific rigor in mice experiments. The Shh*^Cre+/-^* mouse is known to express Cre recombinase under the direction of the *Shh* promoter which is expressed in the epithelium of the lung, bladder, and urogenital organs^39,40^. As such, the Shh*^Cre+/-^* mouse successfully deletes the floxed gene from lung epithelial cells^41^. SPL KO^Epi^ and SPL WT mice were confirmed via genotyping (Fig. 1B). Shh*^Cre+/-^* offspring were utilized for breeding and experiments, as Shh^Cre+/+^ mice were proven to be not viable.

**Figure 1.**
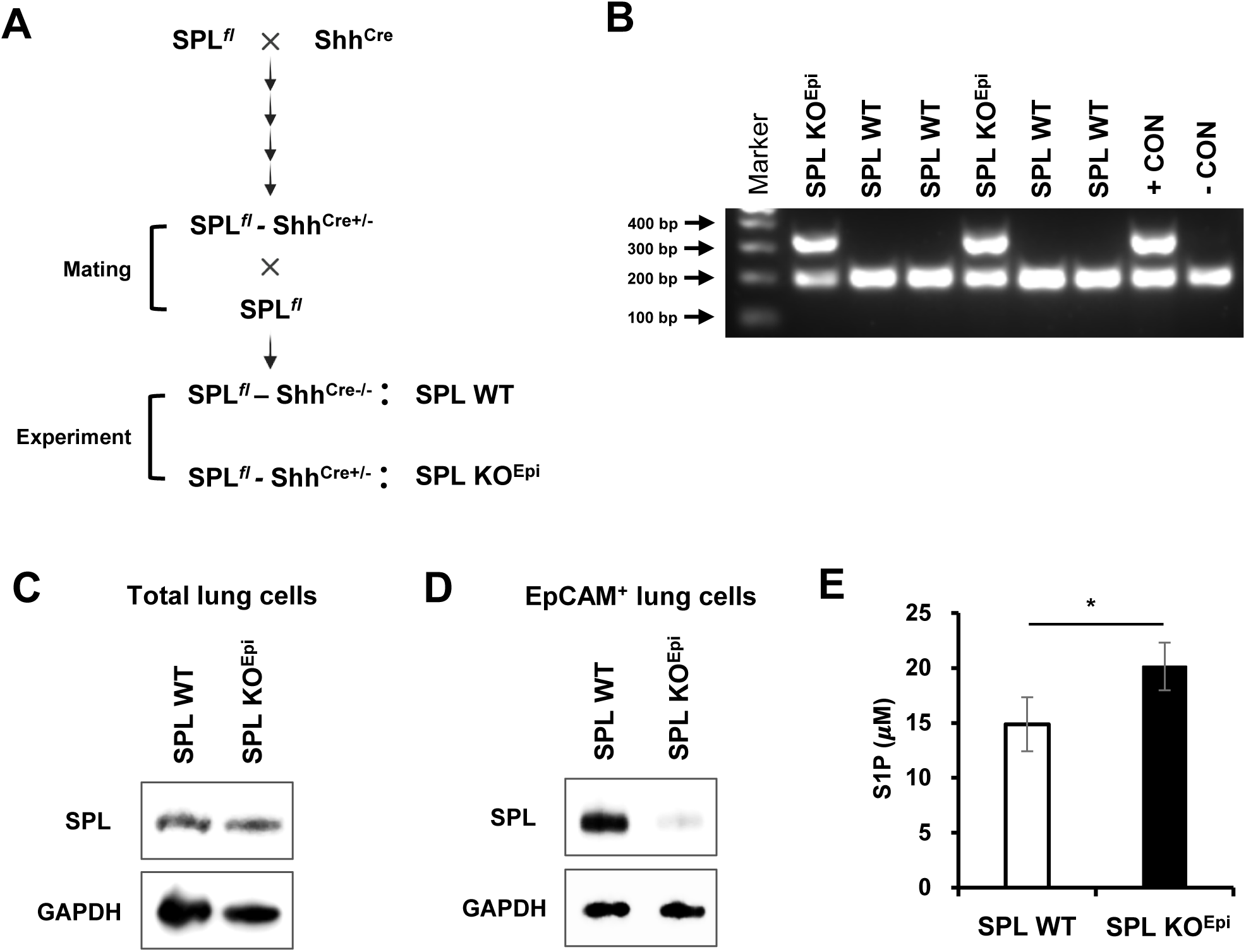
Generation of a conditional SPL KO mouse model and validation of Shh-Cre-mediated SPL deletion from lung epithelial cells. (A) Schematic diagram of the breeding strategy for generating SPL KO^Epi^ mice. SPL*^fl^* mice were crossed with Shh^Cre+/-^ mice to obtain SPL*^fl^* - Shh^Cre+/-^ mice. These heterozygous mice were further backcrossed with SPL*^fl^* mice to generate epithelial-specific knockout (SPL*^fl^* - Shh^Cre+/- ;^ designated as SPL KO^Epi^) mice and their littermate controls (SPL*^fl^* - Shh^Cre-/- ;^ designated as SPL WT) for experiments. (B) Genomic DNA was analyzed to identify SPL WT and SPL KO^Epi^ mice using PCR. The wild-type (WT) alleles and Cre recombinase sequences are indicated by specific band sizes (WT: 200 bp, Cre: 350 bp). Positive (+ CON) and negative (-CON) controls were included for validation. DNA ladder used as size markers (100 - 400 bp) is indicated on the left. (C, D) Immunoblot analysis of SPL expression in isolated lung cells. Western blot analysis was performed on (C) total lung cells and (D) EpCAM^+^ lung epithelial cells purified from SPL WT and SPL KO^Epi^ mice (n=3 pooled mice/group). GAPDH was used as a loading control. The experiment was repeated with similar results. (E) S1P concentrations in serum of SPL WT and SPL KO^Epi^ mice were measured by ELISA. Data are presented as mean ± SEM (n=3 per group). Significance was determined by Student’s *tt*-test. \**p <* 0.05

Knockout of SPL in lung epithelial cells was confirmed by western blot analysis. In total lung cells from SPL KO^Epi^, the expression of SPL protein was not substantially reduced compared to control (Fig. 1C), which is likely due to the low abundance of epithelial cells (∼ 10%) in the total lung cells isolated. However, when epithelial cells were purified from the lung using positive selection of EpCAM, an epithelial cell marker, there was a marked reduction in SPL protein expression (Fig. 1D), confirming epithelial cell-specific removal of SPL. Furthermore, as SPL irreversibly metabolizes S1P, the levels of S1P in serum of SPL WT and SPL KO^Epi^ mice were assessed by ELISA. Levels of serum S1P were modestly but significantly increased in SPL KO^Epi^ mice when compared to S1P levels in SPL WT mice (Fig. 1E), suggesting that SPL enzymatic activity is affected by *Shh* promoter-driven SPL deletion in epithelial cells.

### SPL deficiency in lung epithelial cells results in greater influenza-related morbidity, mortality, and lung viral titers

We previously showed that SPL functions as an antiviral factor during IAV infection in vitro. To evaluate the role of SPL in lung epithelial cells during influenza pathogenesis, 6-7 weeks old SPL WT and SPL KO^Epi^ mice were infected intranasally with 10^2^ plaque forming units (PFU) (0.6 LD_50_) of IAV (PR8) and monitored for their viability over a period of 15 days. SPL KO^Epi^ mice began to succumb to infection at 7 days post infection (dpi). Importantly, IAV infection of SPL KO^Epi^ mice resulted in significantly higher mortality, with only 12% of SPL KO^Epi^ mice surviving, compared to a 63% survival rate of SPL WT mice (Fig. 2A). Further, following IAV infection, the weight loss of SPL WT and SPL KO^Epi^ mice were compared. IAV-infected SPL KO^Epi^ mice showed greater morbidity, losing significantly more body weight than SPL WT mice beginning at 3 dpi. The SPL KO^Epi^ mice lost as much as 28% of their starting body weight and recovered slower than their SPL WT counterparts. Furthermore, only two SPL KO^Epi^ mice remained alive for monitoring at 11 dpi; no SPL KO^Epi^ mice recovered their full initial body weight during the monitoring period of fifteen days. In contrast, surviving SPL WT mice achieved almost full recovery, reaching 98% of their initial weights (Fig. 2B). The results demonstrate that deletion of SPL from lung epithelial cells heightens viral pathogenicity with increased morbidity and mortality.

**Figure 2.**
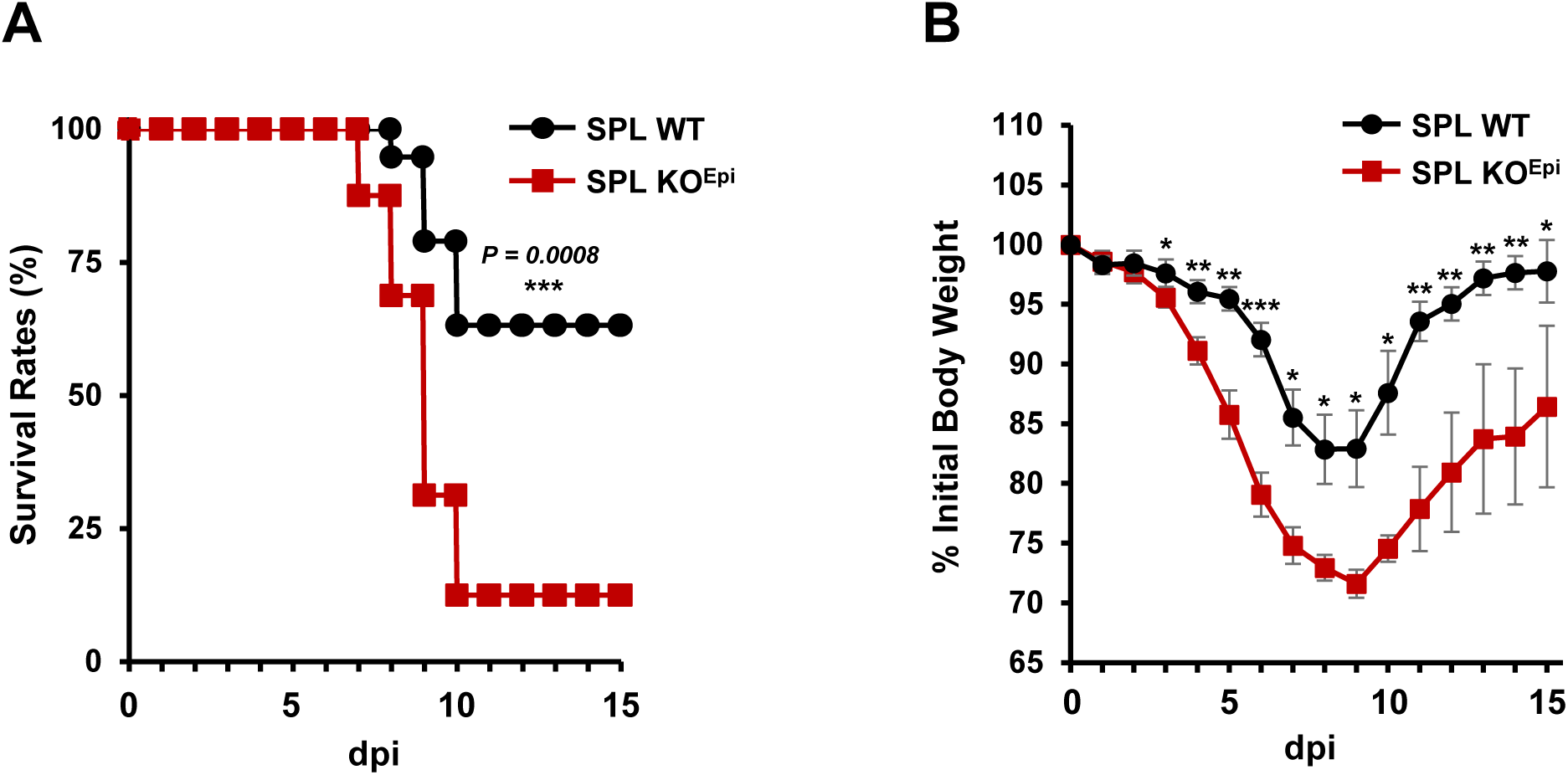
SPL deficiency in lung epithelial cells increased influenza-induced mortality and morbidity. (A) SPL WT (n = 19) and SPL KO^Epi^ (n = 16) mice (6-7 weeks old, mixture of male and female) were infected with 10^2^ PFU of IAV (PR8). Viability of IAV-infected mice was monitored for 15 dpi. Statistical significance was determined using the Kaplan-Meier log-rank test (\*\*\**p* <0.001). Data are combined from two independent experiments. (B) Body weight changes, represented as a percentage of the initial weight, in SPL WT (n=9) and SPL KO^Epi^ (n = 10) mice were recorded for 15 days following IAV infection. The experiment was repeated with similar results. Data are expressed as mean ± SEM. Statistical analysis was performed using Student’s *tt*-test (\**p <* 0.05; \*\**p <* 0.01; \*\*\**p* < 0.001).

As SPL KO^Epi^ mice succumbed to IAV infection at much higher rates than SPL WT mice, we determined if the increased pathogenicity was correlated with robust IAV propagation in the absence of SPL in lung epithelial cells. To this end, IAV titers in lungs of SPL WT vs. SPL KO^Epi^ mice were assessed and compared at various times post infection. While no difference in lung viral titer was observed at 1 dpi between SPL WT and SPL KO^Epi^ mice (Fig. 3A), there was a significant increase in lung viral titer in SPL KO^Epi^ mice at 2 dpi, which was sustained at 6 and 10 dpi (Fig. 3B-D). The results support the antiviral function of SPL in lung epithelial cells in vivo. Furthermore, viral titers from SPL WT mice at 10 dpi show a marked reduction, with some mice displaying no detectable virus, suggesting recovery from infection. However, the SPL KO^Epi^ mice surviving at 10 dpi still harbored substantial amounts of infectious viruses (Fig. 3D), which could explain the delayed recovery in body weights (Fig. 2B).

**Figure 3.**
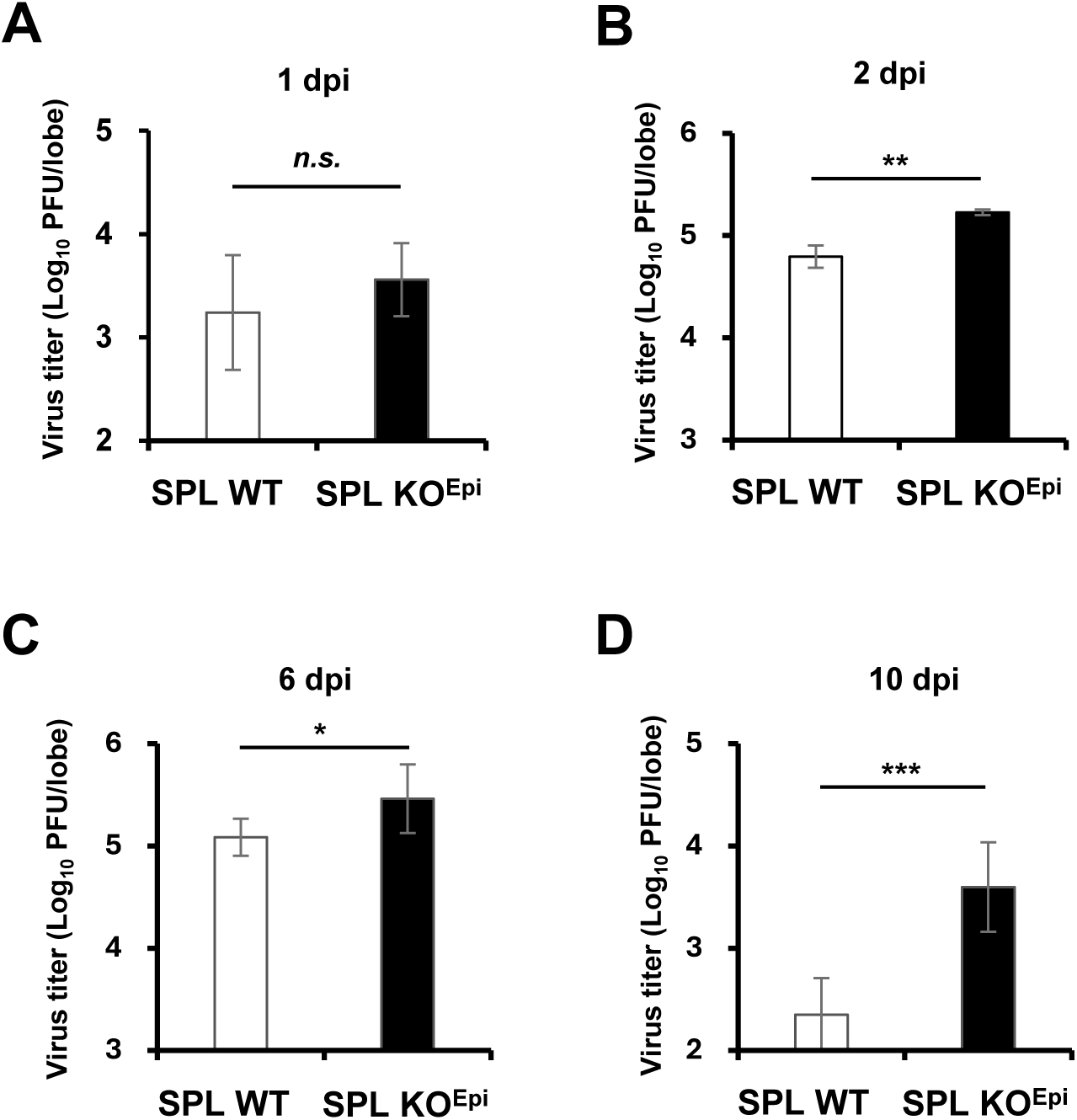
Influenza viral titers increased in the lungs of SPL KO^Epi^ mice compared to SPL WT mice. SPL WT and SPL KO^Epi^ mice were infected with 10^2^ PFU of IAV. Viral titers were determined using lung homogenates of SPL WT and SPL KO^Epi^ mice at (A) 1 dpi (SPL WT, n = 6; SPL KO^Epi^, n = 6), (B) 2 dpi (SPL WT, n = 6; SPL KO^Epi^, n = 4), (C) 6 dpi (SPL WT, n = 6; SPL KO^Epi^, n = 6), and (D) 10 dpi (SPL WT, n = 4; SPL KO^Epi^, n = 4). Data are expressed as Log_10_ PFU per lung lobe. The limit of detection (LOD) is 2 Log_10_ PFU/lobe. Data are expressed as mean ± SD. Statistical analysis was performed using Student’s *tt*-test (\**p <* 0.05; \*\**p <* 0.01; \*\*\**p* < 0.001; *n.s*., not significant).

### Deletion of SPL from epithelial cells leads to enhanced recruitment of immune cells in the lung following IAV infection

Inflammation is known to contribute to influenza pathogenesis^42^. Recruitment of inflammatory immune cells, such as inflammatory monocytes, neutrophils, and macrophages, contribute to enhanced viral pathogenesis^43,44^. To better understand the contribution of immune cell infiltration to increased influenza pathogenicity observed in SPL KO^Epi^ mice, we performed immunophenotyping of lung cells from SPL KO^Epi^ and SPL WT mice at 6 and 10 dpi. Immune cells (CD45^+^) from lung tissue were analyzed via flow cytometry. Ly6C^+^CD11b^+^ inflammatory monocytes, Ly6G^+^CD11b^+^ neutrophils, SiglecF^-^CD11c^+^ dendritic cells (DCs), Ly6G^-^CD11c^+^SiglecF^+^F4/80^+^ alveolar macrophages (AM), and Ly6G^-^SiglecF^-^F4/80^+^ interstitial macrophages (IM) were monitored during infection (Fig. 4A). At both 6 dpi (Fig. 4B-F) and 10 dpi (Fig. 5), we observed a significant increase in all immune cell populations apart from AMs in SPL KO^Epi^ mice when compared to SPL WT mice. Because the SPL KO^Epi^ mice had significantly higher lung viral titer, it is possible that the increased virus within the lung contributed to increased recruitment of immune cells.

**Figure 4.**
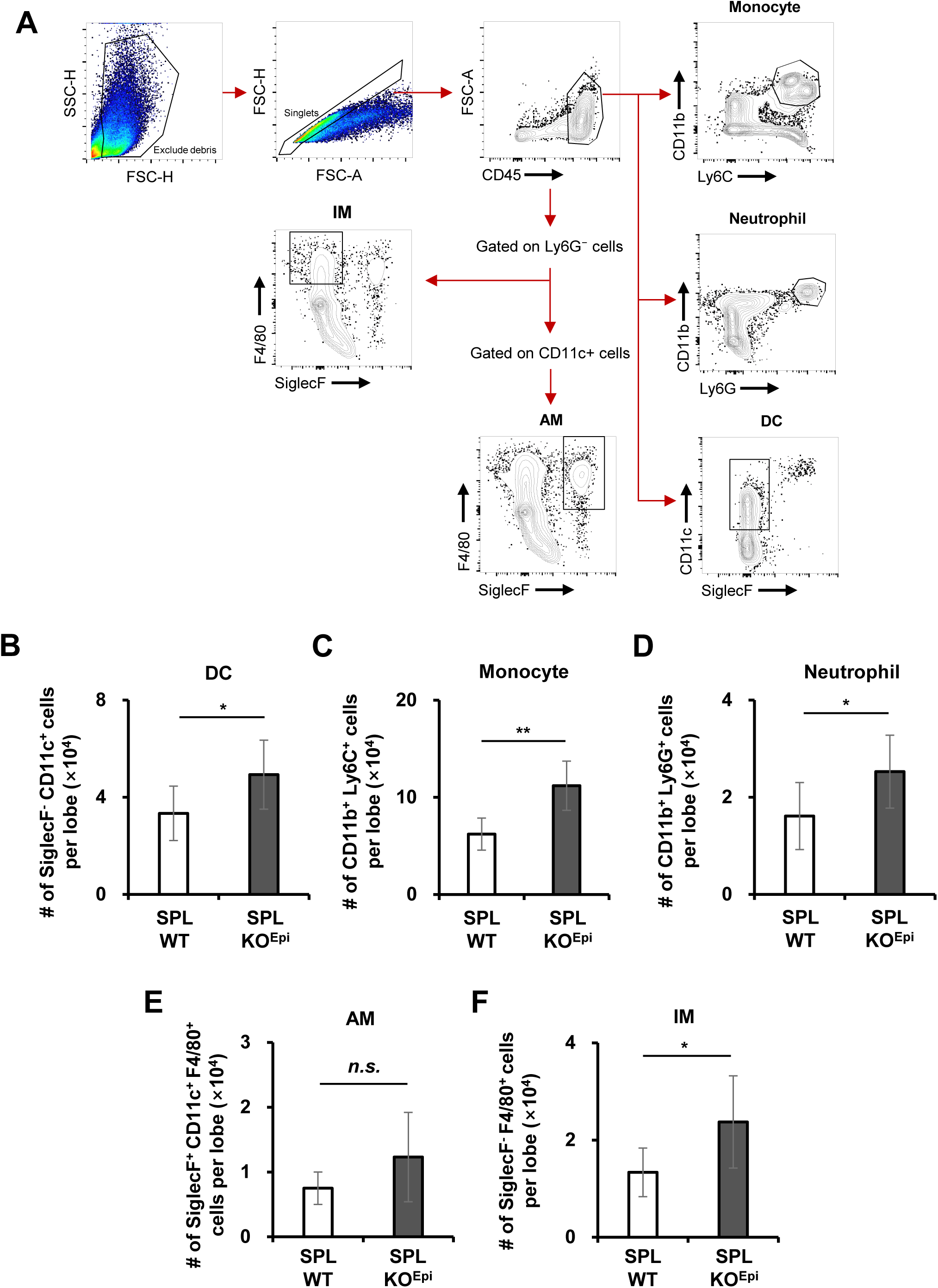
Influenza virus infection increased the accumulation of multiple immune cell types in the lungs of SPL KO^Epi^ mice at 6 dpi. (A) Representative flow cytometry plots illustrating the sequential gating strategy used to identify myeloid cell populations in lung tissues. Following infection with IAV, lung lobes from SPL WT (n=7) and SPL KO^Epi^ (n=6) were collected and dissociated at 6 dpi to prepare single-cell suspensions. Debris was excluded based on forward scatter (FSC-H) and side scatter (SSC-H) profiles, followed by singlet selection using FSC-A and FSC-H. Total leukocytes were gated as CD45^+^ cells. From the CD45^+^ population, monocytes were identified as CD11b^+^Ly6C^+^ cells, neutrophils as CD11b^+^Ly6G^+^ cells, and DCs as CD11c^+^SiglecF^-^ cells. For macrophage subsets, IMs were gated as F4/80^+^SiglecF^-^ cells from the Ly6G^-^population, and AMs were gated as F4/80^+^SiglecF^+^ cells from the Ly6G^-^CD11c^+^ population. (B–F) Absolute cell numbers per lung lobe of (B) DCs, (C) monocytes, (D) neutrophils, (E) AMs, and (F) IMs in SPL WT and SPL KO^Epi^ mice at 6 dpi were measured. Data are expressed as mean ± SD. Statistical analysis was performed using Student’s *tt*-test (\**p <* 0.05; \*\**p <* 0.01; *n.s.*, not significant).

**Figure 5.**
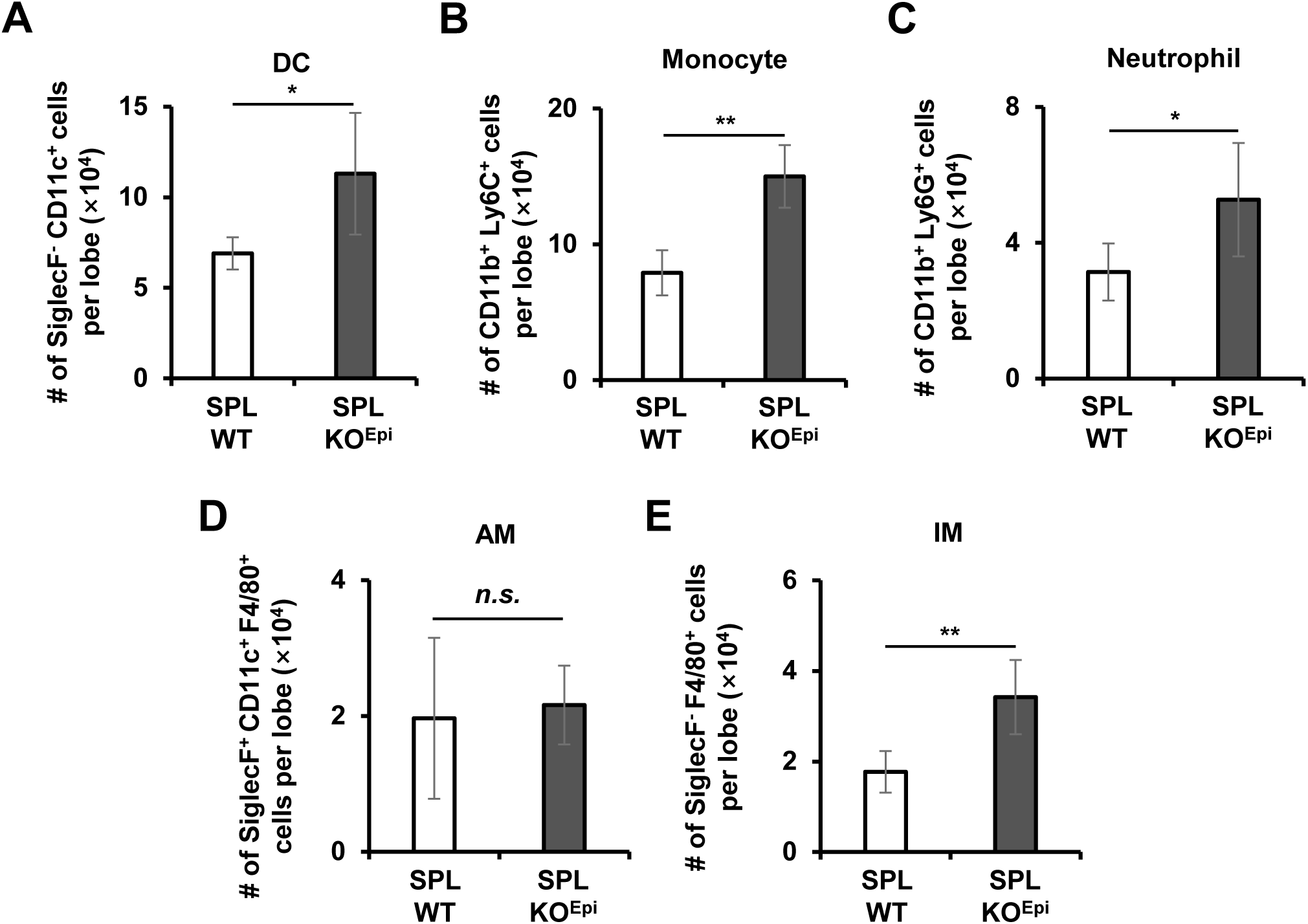
Increases in pulmonary myeloid cell populations in SPL KO^Epi^ mice are sustained at 10 dpi. SPL WT (n=4) and SPL KO^Epi^ (n=5) were infected with IAV. At 10 dpi, flow cytometric analysis was conducted using the experimental and gating strategy shown in Fig. 4A. Absolute cell numbers per lung lobe of (A) DCs (SiglecF^-^CD11c^+^), (B) monocytes (CD11b^+^Ly6C^+^), (C) neutrophils (CD11b^+^Ly6G^+^), (D) AMs (SiglecF^+^CD11c^+^F4/80^+^), and (E) IMs (SiglecF^-^F4/80^+^) were measured. Data are expressed as mean ± SD. Statistical analysis was performed using Student’s t-test (\**p <* 0.05; \*\**p <* 0.01; *n.s*., not significant).

### SPL deficiency in lung epithelial cells induces changes in gene profiles and pathways, including host innate defenses and AT1/AT2 differentiation

To comprehensively elucidate the cellular and molecular landscape within the lung microenvironment during influenza virus infection, we performed scRNA-seq on lung tissues at 2 dpi. Canonical cell-type specific marker genes established in previous murine lung transcriptomic atlases were utilized to identify the cellular subsets^45,46^. Unsupervised clustering and Uniform Manifold Approximation and Projection (UMAP) analysis of total lung cells identified 9 major distinct cell types (Fig. 6A). The identity of each major cell lineage was robustly annotated and validated by evaluating the cell type-specific expression of canonical marker genes using dot plot analysis (Fig. 6B). Since our conditional knockout mouse model targets the deletion of SPL specifically within the lung epithelium, we focused our analysis on the epithelial compartment and performed high-resolution UMAP sub-clustering to dissect the epithelial population in greater detail. This analysis partitioned the lung epithelium into four distinct lineages: AT1, AT2, ciliated, and club cells (Fig. 6C). The annotation of these specialized epithelial subtypes was firmly validated by the cluster-specific expression of canonical lineage markers, as described in the methods section, and presented in both the dot plot (Fig. 6D) and feature plots (Fig. 6E). Intriguingly, compositional analysis revealed a prominent baseline alteration in the epithelial compartments of SPL KO^Epi^ mice (Fig. 6F). Under baseline uninfected conditions, SPL KO^Epi^ mice displayed a markedly elevated proportion of AT1 cells (32.1%) compared to uninfected SPL WT mice (13.4%). This trend persisted following viral infection, with SPL KO^Epi^ mice maintaining a high AT1 cell percentage of 29.8% compared to 12.5% in infected SPL WT mice. These findings demonstrate that epithelial cell-specific SPL deficiency regulates the differentiation of AT2 progenitor cells toward the AT1 lineage.

**Figure 6.**
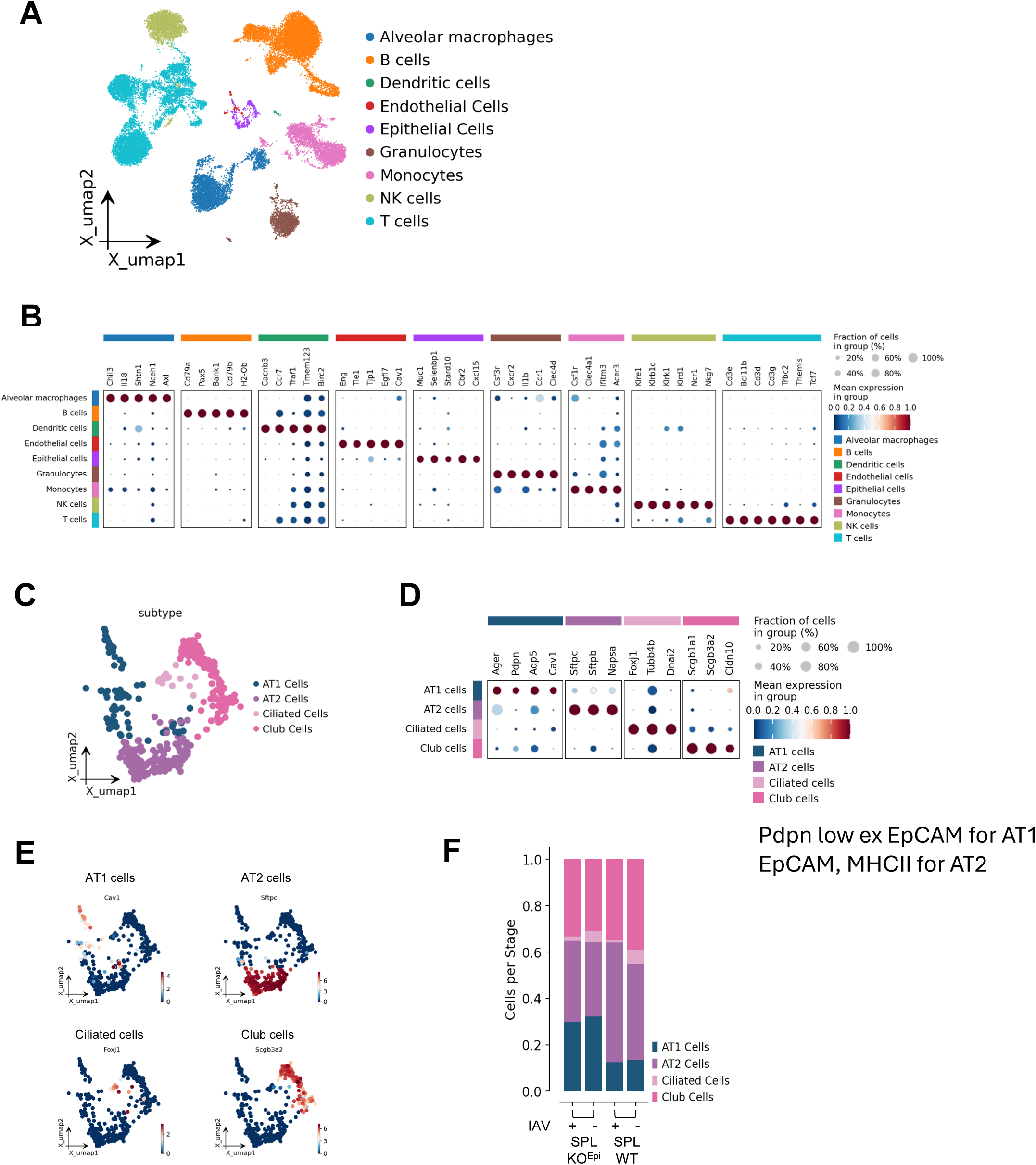
scRNA-seq analysis of lung cells from SPL WT vs. SPL KO^Epi^. SPL WT and SPL KO^Epi^ mice were uninfected or infected with 10^2^ PFU of IAV. At 2 dpi, mouse lung cells were prepared for scRNA-seq analysis. Cells from 3 mice were pooled prior to sequencing. (A) UMAP plot of total lung cells. (B) Marker gene expression across major cell types utilized for cell type annotation. The size of the dot represents the percentage of cells expressing the gene within the cluster and the color intensity represents the average gene expression level within the cluster. (C) UMAP sub-clustering of epithelial cell populations. (D) Expression of lineage-specific marker genes used for annotation of epithelial subpopulations. (E) Visual mapping of localized expression for key lineage-specific genes on the epithelial sub-cluster UMAP. Color scales indicate relative expression intensity. (F) Bar plot illustrating the relative proportion of AT1, AT2, ciliated, and club cells at 2 dpi among the four experimental groups: IAV-infected SPL KO^Epi^ (SPL KO^Epi^ + IAV), uninfected SPL KO^Epi^, IAV-infected SPL WT (SPL WT + IAV), and uninfected SPL WT mice.

To further determine the molecular mechanisms of how epithelial cell-specific SPL deficiency affects lung homeostasis and increases host vulnerability to infection, we analyzed Gene Ontology (GO) biological processes (BP) and gene expression profiles in lung epithelial cells. First, we evaluated the functional differences by examining the significantly enriched pathways in epithelial cells from IAV-infected SPL KO^Epi^ mice compared to IAV-infected SPL WT mice (Fig. 7A). The results revealed that SPL deficiency downregulated the GO BP pathways crucial for host defense, such as host innate immunity, response to type I IFN, and host defense against viruses. This suggests that SPL in lung epithelial cells is crucial for the host’s innate defense against influenza virus infection. Further, genetic pathways of cellular apoptosis regulation, cell stress responses, and inflammation regulation were upregulated, which could be related to the increased virus infectivity and inflammation observed in SPL KO^Epi^ mice. Additionally, pathways of Wnt signaling and response to TGF-β, which are known to regulate AT1/AT2 differentiation^47–49^, were altered in SPL KO^Epi^ mice. We also performed GO analysis of uninfected SPL KO^Epi^ compared to uninfected SPL WT to evaluate the baseline differences before viral infection. This uncovered changes in key pathways, including Wnt signaling regulation and response to TGF-β, suggesting that SPL-mediated regulation of AT1/AT2 programming occurs prior to viral infection (Fig. 7B). Additionally, the top 30 differentially expressed genes (DEGs) between IAV-infected SPL KO^Epi^ and IAV-infected SPL WT epithelial cells were identified (Fig. 7C). Multiple selected genes were reported to regulate host defense mechanisms, including type I IFN responses. For example, *Pbx3* has recently been established as a negative regulator of type I IFN production^50^. Notably, *Pbx3* showed the highest upregulation among the DEGs in epithelial cells of IAV-infected SPL KO^Epi^ mice. *Lyve1*-expressing lymphatic cells function as a critical physical barrier, driven by type I IFN signaling to sequester local viral infection and prevent hematogenous dissemination^51^. *Lyve1* expression was remarkably diminished in epithelial cells from IAV-infected SPL KO^Epi^ compared to IAV-infected SPL WT, suggesting that SPL deficiency compromises this IFN-responsive barrier mechanism (Fig. 7C).

**Figure 7.**
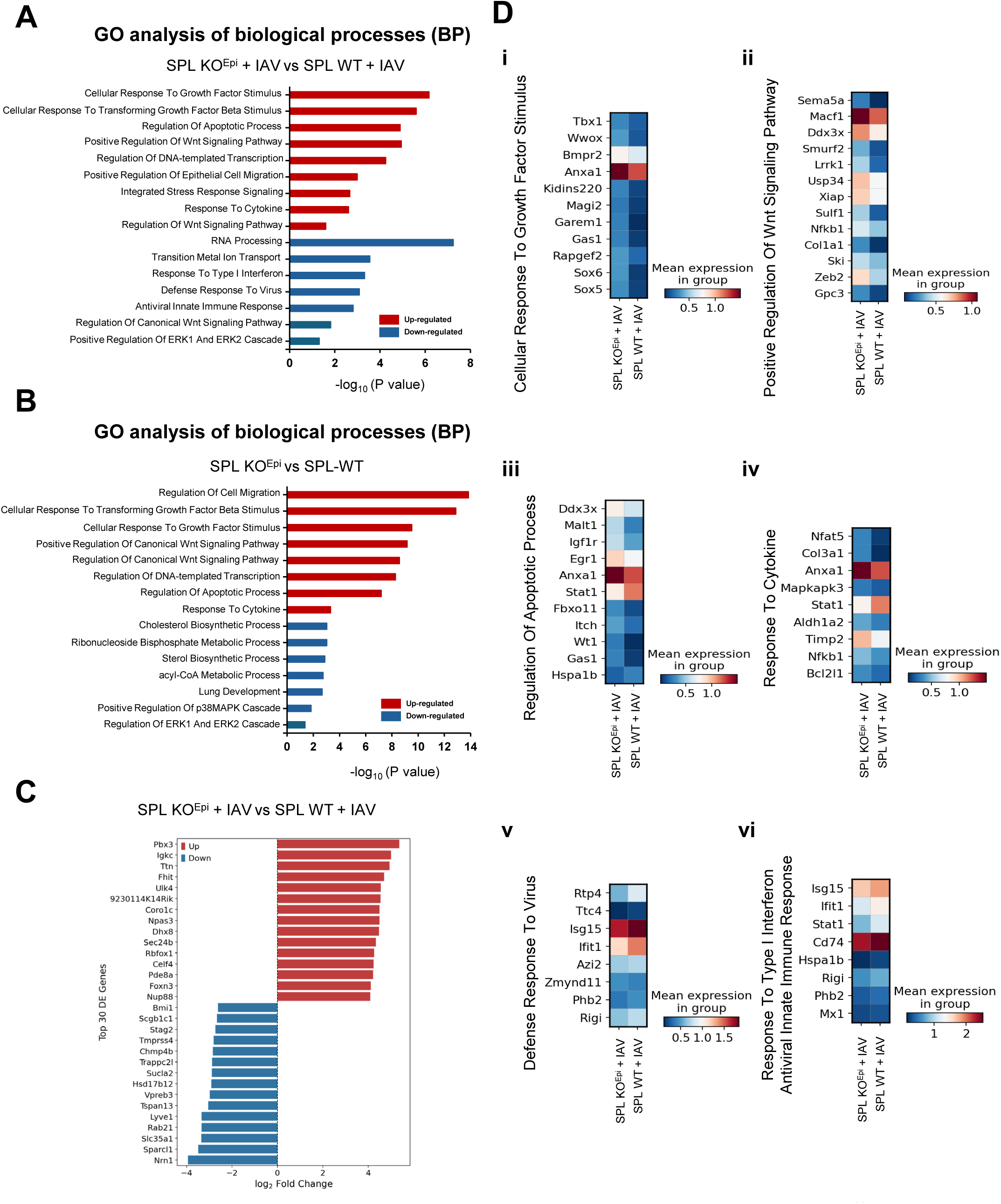
SPL deficiency induces changes in gene profiles and multiple pathways related to host innate defenses, stress responses, and AT1/AT2 differentiation in lung epithelial cells. (A) GO analysis of BP between SPL KO^Epi^ + IAV and SPL WT + IAV mice. Red bars indicate up-regulated processes and blue bars indicate down-regulated processes in SPL KO^Epi^ + IAV compared to SPL WT + IAV, ranked by -log_10_(*P* value). (B) GO analysis of BP between uninfected SPL KO^Epi^ and uninfected SPL WT mice. (C) Top 30 DEGs in SPL KO^Epi^ + IAV versus SPL WT + IAV mice. (D) Heatmaps showing the mean expression levels of representative genes belonging to key altered GO BPs between SPL KO^Epi^ + IAV) and SPL WT + IAV mice: (i) Cellular Response To Growth Factor Stimulus (GO:0071363), (ii) Positive Regulation Of Wnt Signaling Pathway (GO:0030177), (iii) Regulation Of Apoptotic Process (GO:0042981), (iv) Response To Cytokine (GO:0034097), (v) Defense Response To Virus (GO:0051607), and (vi) Response To Type I Interferon (GO:0034340) and Antiviral Innate Immune Response (GO:0140374).

In-depth analysis was performed to determine the expression of specific DEGs within identified GO categories (Fig. 7D). We identified expression shifts in genes critical for “Cellular Response To Growth Factor Stimulus” (Fig. 7D-i) and “Positive Regulation Of Wnt Signaling Pathway” (Fig. 7D-ii). Key components of the Wnt pathway, such as *Ddx3x*, *Smurf2*, and *Zeb2*, showed clear expression increases in the SPL KO^Epi^ epithelial population. Because carefully controlled Wnt signaling is required to maintain progenitor cells and repair the lung epithelium^48,52^, these changes suggest that SPL plays an important role in controlling Wnt-driven epithelial cell differentiation. Further, genes involved in the “Regulation Of Apoptotic Process” (Fig. 7D-iii) were significantly increased, which may reflect the increased virus infection or a role of SPL in regulation of cellular apoptosis. Importantly, genes related to “Defense Response To Virus” (Fig. 7D-v) and “Response To Type I Interferon / Antiviral Innate Immune Response” (Fig. 7D-vi), including key antiviral genes such as *Isg15, Ifit1*, and *Cd74*^53–56^, showed a clear decrease in epithelial cells of IAV-infected SPL KO^Epi^ compared to IAV-infected SPL WT. Consequently, the impaired antiviral barrier likely caused a more severe and uncontrolled viral infection within the SPL-deficient lung epithelial cells.

## Discussion

The newly created mouse model with conditional deletion of SPL from lung epithelial cells allowed us to determine the role of SPL during influenza in vivo. SPL deficiency increased influenza viral propagation in the lungs and consequently worsened influenza-induced weight loss and mortality, demonstrating the antiviral function of SPL in respiratory epithelium during influenza.

Influenza viral titer began to increase between 1 and 2 dpi in the absence of SPL (Fig. 3), which may have resulted in the significant weight loss of SPL KO^Epi^ mice compared to control mice initiated between 2 and 3 dpi (Fig. 2B). The heightened viral burden in the SPL KO^Epi^ mouse likely alarmed the host immune system and triggered an increased activation of immune cells that transmit signals to induce the infiltration of inflammatory immune cells, such as neutrophils and inflammatory monocytes (Fig. 4 and 5). While monocytes, IMs, DCs, and neutrophils increased, the number of AMs did not significantly change in SPL KO^Epi^ mice compared to SPL WT controls upon infection. Lung resident AMs were previously reported to be depleted upon influenza virus infection^57^. Thus, it is possible that the increased infectivity of SPL KO^Epi^ mice may have more intensively removed AMs than in the control WT mice, compensating for the increase in AMs observed in SPL KO^Epi^ mice. As excessive inflammatory responses are injurious to the lungs, the continuously increased accumulation of inflammatory immune cells in the lungs of SPL KO^Epi^ mice could have contributed to influenza pathogenesis and the subsequent increased severity.

AT1 and AT2 cells are the primary epithelial cells making up the alveoli. AT1 cells are dedicated to gas exchange while AT2 cells secrete surfactants and act as stem-like progenitor cells that differentiate into AT1 cells to repair lung injury^58^. The scRNA-seq results indicate that SPL deficiency by itself increases the proportion of AT1/AT2 with alteration of genetic pathways important for AT1/AT2 differentiation, such as Wnt signaling and TGF-β responses. This suggests that SPL in lung epithelial cells regulates AT1/AT2 programming. This finding agrees with the previous report that sphingosine kinase 1 in endothelial cells produces S1P and regulates AT1/AT2 differentiation^59^. Thus, S1P increased by SPL deficiency could have promoted AT1 differentiation from AT2 cells. However, since S1P could act intracellularly as well as extracellularly, whether there is a specific epithelial cell type important for SPL regulation of AT1/AT2 differentiation remains an open question. More importantly, it is unclear if the increased AT1 cells affected influenza pathogenicity in SPL KO^Epi^ mice. Further, we cannot exclude the possible contribution of deficiency of S1P catabolic products, i.e., hexadecenal and phosphoethanolamine, to the observed phenotype. While both AT1 and AT2 cells can be productively infected with influenza virus, it was reported that influenza virus exhibits a higher infectivity and tropism toward AT1 cells over AT2 cells^60^. Thus, it is possible that prior to viral infection, alteration in lung alveolar homeostasis through an increased AT1 cell population potentially increased susceptibility to influenza virus infections, exacerbating respiratory distress. Alternatively, given that AT1 cells are important for gas exchange and lung repair, increased AT1-mediated lung function could represent a beneficial effect of SPL deficiency. However, it may not be able to compensate for the worsened influenza burden caused by the loss of SPL’s direct antiviral activity. Additional animal models should be established to address these present scientific questions.

The gene signature pathway analysis uncovered that SPL in lung epithelial cells is important for the host defense and innate immunity, such as type I IFN responses. The results obtained in vivo are consistent with our previous in vitro findings supporting pro-type I IFN functions of SPL observed in A549 and HEK cell lines. The scRNA-seq analysis of top DEGs further reveals specific genes associated with these pathways. While some IFN-stimulated genes (ISGs), such as *Isg15* and *Ifit1*, are already well reported to inhibit influenza virus replication^53,56^, others, such as *Pbx3*, *CD74*, and *Phb2,* remain to be investigated during influenza.

Sphingolipid-metabolizing enzymes have been investigated during influenza. However, most of the studies have been limited to the use of inhibitors or in vitro tissue culture systems. The development of conditional KO animal models should help to further define their cell type specific functions in vivo during influenza and identify new host factors in the pathways of virus-host interplay that better control viral pathogenicity.

## Methods

### Mice

All mice used in this study were generated on a C57BL/6 background (The Jackson Laboratory). *Sgpl1^fl/fl^* (SPL*^fl^*) mice were crossed with *Shh^Cre^* mice (The Jackson Laboratory) for at least four generations to generate SPL*^fl^*-Shh*^Cre+/-^* mice. SPL*^fl^*-Shh*^Cre+/-^*were then backcrossed with SPL*^fl^* to generate the SPL KO^Epi^ and littermate SPL WT mice for experiments. Mice were housed with a maximum of five sex-matched mice per cage and had ad libitum access to food and water. Six- to eight-weeks old male or female mice were used for experiments. During experiments, mice were monitored daily. Humane euthanasia via CO_2_ inhalation (4L/min flow rate) in home cages (75 inch^2^) was conducted for any mouse that was found moribund. Mice that lost 25% or more of their initial body weight were monitored for an additional 24 hours. If no improvement was observed, the mouse was euthanized. At the end of the experimental period, all mice were euthanized. Mice were bred and maintained in a closed breeding facility according to institutional guidelines and animal protocols approved by the Animal Care and Use Committee of University of Missouri-Columbia.

### Virus and infections

Mice were infected intranasally with 100 PFU of influenza A/Puerto Rico/8/34 (H1N1) virus (PR8) (originally provided by Adolfo Garcia-Sastre, Mount Sinai School of Medicine). For viral infection, mice were anesthetized using a SomnoFlo® vaporization system with 5% isoflurane administered at a high flow rate for 10–15 s. 40 µl of PBS containing virus or vehicle was administered to anesthetized mice intranasally^20,61^. Following infection, mice were monitored to ensure full recovery from anesthesia.

### Genotyping

Ear punch samples from mice were collected and used for DNA extraction. Briefly, samples were incubated in a solution containing proteinase K (GoldBio) and digested overnight in a water bath. DNA was extracted using ethanol and finally resuspended in 1X TE buffer. DNA was then used for polymerase chain reaction (PCR) to detect the presence of *SPL* and *Shh-Cre*. Two different primers were used for the detection of *SPL*: Primer 1 - 5’ GAA ATT GAG CAT ATC CGT TC 3’; Primer 2 - 5’ GTT CTG GAT GGA GTT TA 3’. Three different primers were used for detection of *Shh-Cre*: Primer 1 – 5’ GGT GCG CTC CTG GAC GTA 3’; Primer 2 – 5’ GGG ACA GCT CAC AAG TCC TC 3’; Primer 3 – 5’ CTC GGC TAC GTT GGG AAT AA 3’. All primers were obtained from IDT. DNA was mixed with a PCR master mix containing 2x Go Taq polymerase (Promega), then loaded into a Bio-Rad T100 Therma Cycler to carry out the reaction. Samples were then loaded and run on a 2% agarose gel, followed by imaging with an Odyssey Fc (LI-COR) machine and analysis using Image Studio V5.2 (LI-COR).

### Lung collection and cell isolation

Lungs were collected from mice after euthanasia via CO_2_ inhalation and kept on ice until processed. Lung homogenates for western blot, flow cytometry, and scRNA-seq were prepared using a gentleMACS dissociator (Miltenyi Biotec) according to manufacturer directions. Briefly, lungs were dissociated and incubated in media containing collagenase D (Sigma-Aldrich) and DNAse I (Sigma-Aldrich). Cells were pelleted and filtered using a 100 µm strainer. Red blood cells were lysed using 1X RBC lysis buffer (Sigma Aldrich). Debris was removed from cells for western blot and flow cytometry by Percoll® (Sigma Aldrich) treatment. Cells for western blot were enriched for EpCAM-positive epithelial cells using an epithelial cell isolation kit (Stemcell Technologies) and then suspended in 1X sample buffer containing β-mercaptoethanol and used for further analysis. Cells for flow cytometry were suspended in PBS containing 1% FBS and ready for staining. Lungs for viral titration were incubated in PBS supplemented with 1 µg/ml TPCK-treated trypsin (Sigma-Aldrich), then transferred to 2 ml screw-top vials containing 1.0 mm zirconia/silica beads. Lungs were homogenized using a Mini-BeadBeater-16 (BioSpec Products) and subjected to two cycles of shaking for 30 s each. Shaken tubes were incubated on ice for 2 minutes between rounds of shaking. The resulting homogenate was collected and ready to be used for plaque assay.

### Western blotting

Protein lysates were boiled on a heat block then vortexed. Lysates were loaded onto 10% sodium dodecyl sulfate-polyacrylamide gel electrophoresis (SDS-PAGE) gel and run. Proteins were transferred to a nitrocellulose membrane, blocked with 5% non-fat dry milk for 1 hour at room temperature, and incubated with primary antibody against SPL (Invitrogen) and GAPDH (Cell Signaling Technology) overnight. Membranes were then incubated with HRP-linked anti-rabbit antibody (Cell Signaling Technology) for 1 hour^35,62,63^. Signals were detected using an enhanced chemiluminescence substrate (Thermo Scientific) and imaged with an Odyssey Fc (LI-COR) machine and analyzed with Image Studio V5.2 (LI-COR).

### S1P ELISA

S1P concentrations in serum of SPL WT and SPL KO^Epi^ mice were assessed using a competitive S1P ELISA kit (Echelon Biosciences) according to the manufacturer’s instructions. Briefly, serum samples were diluted 1:10 in the provided delipidized serum. The S1P standards were prepared utilizing serial dilutions (2, 1, 0.5, 0.25, 0.125, 0.0625, and 0 µM) in delipidized serum to generate the standard curve. Both serum samples and standards were then combined with biotinylated anti-S1P antibody before being transferred to blocked, S1P-coated microplate wells and incubated for 1 hour at room temperature. Following incubation, HRP was added to the plate and incubated for 1 hour. Tetramethylbenzidine (TMB) substrate was added and incubated for 30 minutes at room temperature, followed by the addition of 1 N sulfuric acid to halt the reaction^64,65^. Absorbance was measured at 450 nm using a BioTek Epoch microplate reader (Agilent Technologies). A standard curve was generated using GraphPad Prism (V10.2.0), and S1P concentrations were interpolated and corrected for the 1:10 sample dilution.

### Plaque assay

Virus titration was conducted via plaque assay as previously reported^20,36,62,63^. Briefly, Madin-Darby canine kidney (MDCK) cells at least 90% confluent were washed with PBS and infected with 10-fold serial dilutions of virus-containing lung homogenate, as previously described. Virus was diluted in PBS containing 0.3% bovine serum albumin (BSA) and supplemented with 1% GlutaMax (Gibco). Cells were incubated with virus at 37°C and 5% CO_2_ for one hour. Virus was removed and infected cells were overlaid with a PBS mixture containing 0.6% SeaKem LE agarose (Lonza), 2x L15 (Invitrogen), 0.3% BSA, and 1 µg/ml TPCK-treated trypsin (Sigma-Aldrich). After an incubation of 3 days at 37°C and 5% CO_2_, cells were fixed for at least two hours with 4% formaldehyde. Overlays were removed and cells were stained using methanol containing 2% crystal violet to visualize plaques and quantify viral titer.

### Flow cytometry

Cells isolated from lung tissue as described above were counted and at least 5×10^5^ cells/well were seeded in a 96-well V-bottom plate for staining. Cells were pelleted and resuspended in antibody cocktail. All antibodies used were purchased from BioLegend: CD45-BV785, CD45-APC Fire 810, CD11b-PerCP-Cy5.5, Ly6C-FITC, Ly6G-APC, CD11c-PE-Cy7, SiglecF-BV421, SiglecF-PE-Dazzle 594, and F4/80-PE-Dazzle 594. Cells were incubated with antibodies in the dark for 30 minutes at 4°C. After incubation, cells were washed twice with PBS containing 1% FBS. Supernatant was discarded and the cells were fixed with 2% paraformaldehyde solution^66,67^. Samples were analyzed using the Cytek Aurora (Cytek Biosciences) flow cytometer and analyzed with FlowJo (V10.10.1).

### Single-cell RNA sequencing

#### Cell processing

Cells prepared from lung homogenate as described above were subjected to an additional round of filtration using a 100 µm strainer and finally suspended in PBS containing 1% FBS. To minimize contamination from dead cells and ambient RNA released during tissue dissociation, single-cell suspensions were stained with DAPI and DAPI-positive dead cells were excluded by flow cytometric cell sorting. Library preparation and sequencing were conducted through the University of Missouri-Columbia Genomics Technology Core facility. Gene libraries were generated using the Chromium Next GEM Single Cell 3’ Kit (V3.1, 10X Genomics) according to manufacturer’s instructions. Cell viability was further evaluated using acridine orange and propidium iodide staining (Invitrogen). Emulsions with gel beads were prepared, followed by reverse transcription, cDNA amplification, and library construction according to 10X Genomics protocol. Sequencing was conducted using an Illumina NovaSeq X Plus with a depth of 50,000 reads per cell, sequencing 10,000 cells per sample^67^.

#### Single-cell RNA-seq data processing and analysis

The scRNA-seq data from mouse lung samples were processed using the Cell Ranger toolkit (v9.0.1), with reads aligned to the GRCm39 mouse reference genome. Reads that were uniquely aligned, not marked as PCR duplicates, and assigned to valid cell barcodes and unique molecular identifiers (UMIs) were used to generate the initial 3′ gene-by-cell count matrix, which contained 37,099 cells. To reduce ambient RNA contamination and background-associated artifacts, the initial count matrix was processed using CellBender (v0.3.2)^68^. Cells were retained if they passed a false discovery rate (FDR) threshold of <0.01 after Benjamini–Hochberg correction. Additional quality-control filters were applied to remove low-quality cells, including cells with ≤1,000 total UMIs, ≤500 detected genes, or ≥15% mitochondrial transcript content. Potential doublets were further removed using a cluster-level filtering strategy. Doublet scores were calculated for each cell using DoubletDetection (v4.3)^69,70^. For each sample, cells were clustered by selecting the top 3,000 highly variable genes, performing principal component analysis (PCA), and retaining the top 50 principal components. A k-nearest-neighbor graph was then constructed using the pp.neighbors function in Scanpy (v1.11.0)^71^, followed by Louvain clustering. Within each cluster, doublet scores were median-centered and scaled by the median absolute deviation (MAD), and cells or clusters with elevated scaled doublet scores were removed as likely doublets. After quality control and doublet removal, 25,814 high-confidence cells from four biological samples representing 12 mice were retained for downstream analysis. For each cell, both raw UMI counts and log₂-transformed normalized expression values were computed.

#### Batch correction and data integration

To account for inter-sample variability and batch-associated confounders, cells were integrated using scvi-tools (v0.20.0)^72^. scvi-tools uses scVI, a probabilistic generative model built on variational autoencoders, to project single-cell expression data into an integrated latent space that corrects technical batch effects without removing relevant biological differences.

#### Clustering, cell-type annotation, and differential expression

For downstream clustering, 3,000 highly variable genes were selected, and PCA was performed to extract the top 50 principal components. A k-nearest-neighbor graph was constructed using the pp.neighbors function in Scanpy^71^, and clustering was performed using the Leiden algorithm at a resolution of 0.5. This analysis identified 25 transcriptionally distinct clusters, which were visualized using UMAP^73^. Initial cell-type annotation was performed using SingleR (v2.10.0)^73^ with the MouseRNAseqData reference from Celldex (v1.18.0). These annotations were further refined by calculating gene signature scores with the Scanpy^71^ tl.score_genes function using curated canonical marker genes. AMs were characterized by high expression of *Chil3*, *Il8*, *Shtn1*, *Nceh1*, and *Axl*. B cells were identified by expression of *Cd79a*, *Pax5*, *Bank1*, *Cd79b*, and *H2-Ob*. DCs were defined by expression of *Cacnb3*, *Ccr7*, *Traf1*, *Tmem123*, and *Birc2*. Endothelial cells were annotated based on expression of *Eng*, *Tie1*, *Tjp1*, *Egfl7*, and *Cav1*, whereas epithelial cells were marked by *Muc1*, *Selenbp1*, *Stard10*, *Cbr2*, and *Cxcl15*. Granulocytes showed elevated expression of *Csf3r*, *Cxcr2*, *Il1b*, *Ccr1*, and *Clec4d*. Monocytes were characterized by expression of *Csf1r*, *Clec4a1*, *Ifitm3*, and *Acer3*. NK cells were identified by expression of *Klre1*, *Klrb1c*, *Klrk1*, *Klrkd1*, *Ncr1*, and *Nkg7*. T cells were annotated based on expression of *Cd3e*, *Bcl11b*, *Cd3d*, *Cd3g*, *Trbc2*, *Themis*, and *Tcf7*. To investigate transcriptional heterogeneity within the epithelial compartment, a second round of clustering was performed using only cells annotated as epithelial cells. Gene module scores were calculated for epithelial subpopulation classification based on curated marker sets. AT1 cells were defined by elevated expressions of *Ager*, *Pdpn*, *Aqp5*, and *Cav1*. AT2 cells were enriched for *Sftpc*, *Sftpb*, and *Napsa*. Ciliated cells showed high expression of *Foxj1*, *Tubb4b*, and *Dnai2*, whereas club cells were characterized by expression of *Scgb1a1*, *Scgb3a2*, and *Cldn10*. DEGs were identified using the Scanpy^71^ tl.rank_genes_groups function with two-sided t-test, followed by Benjamini–Hochberg correction for multiple testing. Genes with an adjusted P value <0.05 and a mean log₂ fold change >0.5 were considered statistically significant. The complete scRNA-seq dataset has been deposited in the GEO database under accession number GSE337022.

#### Functional enrichment analysis

To assess the biological relevance of cluster-specific marker genes, gene set enrichment analysis was performed using GSEApy (v1.1.8)^74^. Differential expression results comparing IAV-infected SPL KO^Epi^ samples with IAV-infected SPL WT samples were used to define upregulated and downregulated gene sets. These gene sets were queried against curated databases, including Enrichr^75^ and GO BP collections, to identify pathways and biological processes associated with the epithelial-cell transcriptional response to treatment.

### Statistical analysis

Error bars in all figures indicate either the mean values ± standard deviations (SD) or mean values ± standard errors of the mean (SEM), as specified in the respective figure legends. Statistical significance was determined using Student’s *t*-test (*p* < 0.05). For animal mortality and survival data, statistical significance between groups was evaluated using the log-rank (Mantel-Cox) test.

## Acknowledgments

We thank Regina Wamsley and Matthew Bogan for assistance with mouse maintenance and genotyping. We acknowledge the Office of Animal Resources at the University of Missouri for animal care. We also thank the Cell and Immunobiology Core and Genomics Technology Core at the University of Missouri for technical support. This work was supported by NIH/NIAID R01AI153076 and R01AI162631 (BH) and NIH/NHLBI HL171786 (JDS).

## Data Availability Statement

The dataset used for scRNA-seq in the present study can be found in the GEO database (GSE337022). Data used in preparing this article will be released upon request to the corresponding author.

